# Shifts in isoform usage underlie transcriptional differences in regulatory T cells in type 1 diabetes

**DOI:** 10.1101/2022.09.07.506965

**Authors:** Jeremy R. B. Newman, S. Alice Long, Cate Speake, Carla J. Greenbaum, Karen Cerosaletti, Stephen S. Rich, Suna Onengut-Gumuscu, Lauren M. McIntyre, Jane H. Buckner, Patrick Concannon

## Abstract

Genome-wide association studies have identified numerous loci with allelic associations to Type 1 Diabetes (T1D) risk. Most disease-associated variants are enriched in regulatory sequences active in lymphoid cell types, suggesting that lymphocyte gene expression is altered in T1D. We assayed gene expression between T1D cases and healthy controls in two autoimmunity-relevant lymphocyte cell types, memory CD4+/CD25+ T-regulatory cells (Treg) and memory CD4+/CD25- T-cells, using a splicing event-based approach to characterize tissue-specific transcriptomes. Limited differences in isoform usage between T1D cases and controls were observed in memory CD4+/CD25- T-cells. In Tregs, 553 genes demonstrated differences in isoform usage between cases and controls, particularly RNA recognition and splicing factor genes. Many of these genes are regulated by the variable inclusion of exons that can trigger nonsense mediated decay. Our results suggest that dysregulation of gene expression, through shifts in alternative splicing in Tregs, contributes to T1D etiology.

## Introduction

Type 1 diabetes (T1D) is an autoimmune disease arising from the T cell-mediated destruction of the insulinogenic pancreatic β cells, resulting in complete dependence on exogenous insulin to maintain glucose homeostasis ^1,2^. A substantial genetic contribution to the disorder is well-established ^3–5^. Up to half of the genetic risk for T1D is attributed to the human leukocyte antigen (HLA) gene cluster on chromosome 6 ^6–8^. There are also more than 90 non-HLA chromosomal regions for which significant evidence of association with T1D exists ^9^. Fine mapping with the ImmunoChip of non-HLA regions associated with T1D combined with Bayesian inference has established a set of highly credible, putatively causative SNPs at many of these loci ^10^. However, only a few of these credible causative variants are located in the coding regions of genes. Interrogation of 15 chromatin states across 127 tissues at the chromosomal positions of these credible SNPs revealed a strong enrichment for transcriptional enhancer sequences active in lymphocytes and other immune-relevant tissues ^10^, suggesting that changes in transcriptional regulation may be the mode of action for many of these T1D risk loci. These results are consistent with the hypothesis that most genetic variants that contribute towards disease risk are located in non-coding regions of the genome and modify gene regulation rather than impacting directly on protein function.

Dysregulation of transcription has been implicated in many human diseases ^11–19^, and can take the form of changes in overall transcriptional abundance of specific genes between affected and unaffected individuals or through alternative splicing that leads to alterations in transcript production, rather than gene usage. Alternative splicing of several genes in lymphocytes has been shown to be modified by T1D-associated risk variants located in or near those genes ^20–24^. In *UBASH3A*, a rare alternate allele (G) at rs56058322 in intron 9 confers protection against T1D and favors the production of a truncated, intron-retaining isoform ^25,26^. In *PTPN22*, a rare missense alternate allele (G) at rs56048322 in exon 18 is associated with T1D risk and results in the expression of two novel transcripts ^27,28^.

In this study we systematically evaluate transcript events in CD4+ T cells and whether lymphoid transcriptional dysregulation, in the form of alternative splicing, contributes towards T1D pathology. We broadly examine gene expression, splicing and isoform usage in subpopulations of memory CD4+ T cells, fractionated on their expression of CD25 (memory CD4^+^/CD25^+^ Tregs, memory CD4^+^/CD25^-^ T cells) in order to elucidate transcriptional mechanisms underlying T1D in these cell types.

## Results

### Defining cell type-specific reduced reference transcriptomes

The majority of detected transcriptional events were observed in both T1D cases and unaffected controls (Figure 1A), and most events could be annotated to known transcripts (i.e., exon fragments and previously reported junctions), across the interrogated cell types. However, cell type specific differences were observed among unannotated events, i.e., previously undescribed junction and exon-intron border junctions. Unannotated events were more likely to be group specific in Tregs than in memory CD4^+^/CD25^-^ T cells (Figure 1A), and among Tregs there were more unannotated events detected in T1D cases than controls.

**Figure 1.**
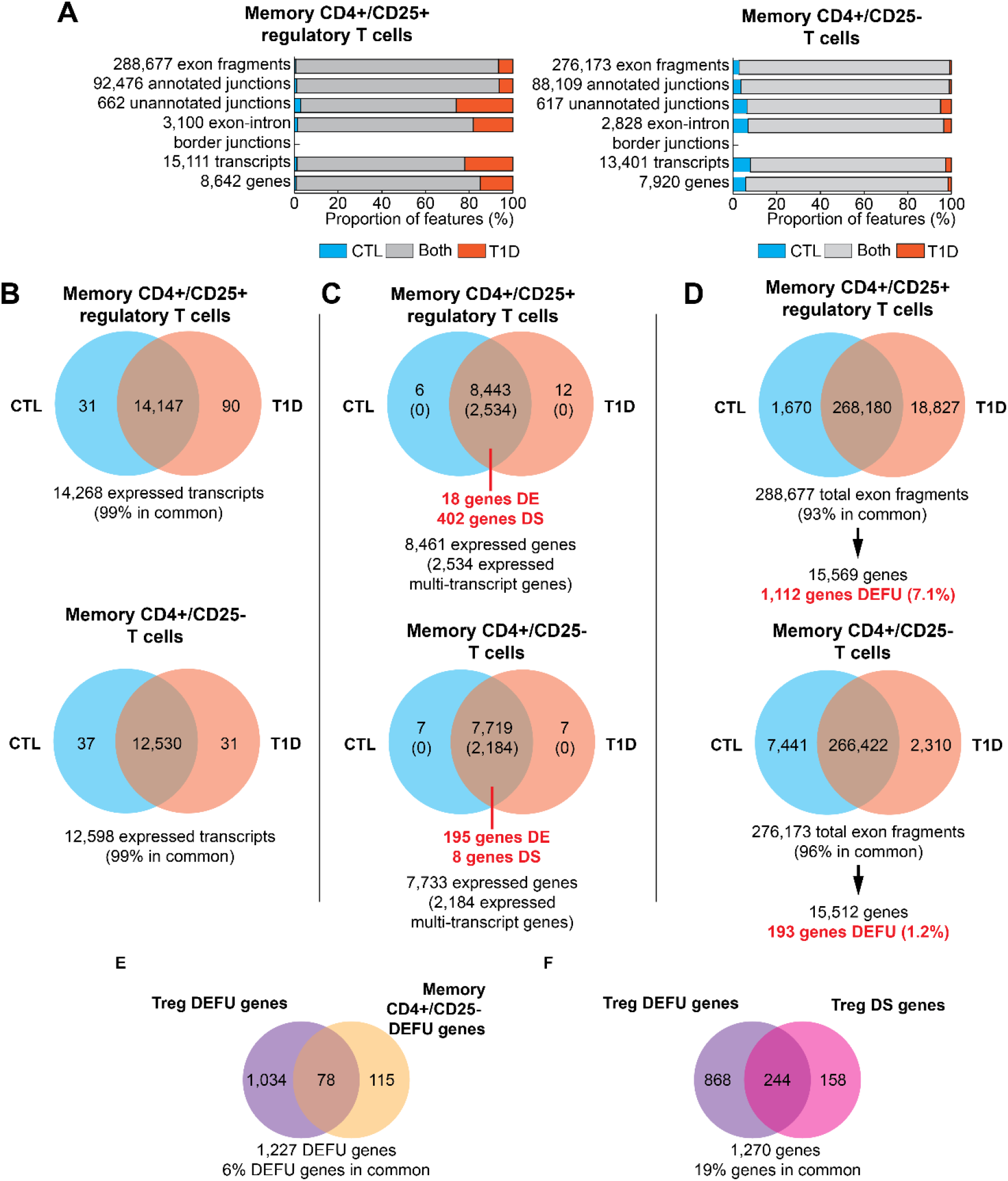
Summary of gene expression and splicing analysis. (A) Transcriptional events – exon fragments, exon-exon junctions, and exon-intron border junctions – detected at APN≥2 in T1D cases (orange), unaffected controls (CTL; blue), and in both (grey) in memory CD4^+^/CD25^+^ Tregs, memory CD4^+^/CD25^-^ T cells, and total memory CD4^+^ T cells, and the resulting reduced transcript sets with all associated events detected. (B) Transcripts detected at TPM>0 in Tregs and memory CD4^+^/CD25^-^ T cells at TPM>0. (C) Genes detected at TPM>0. Numbers of detected multi-transcript genes are presented in parentheses. The number of significantly differentially expressed (DE) genes and differentially spliced (DS) genes (FDR P<0.05) for each cell type is displayed in red text. (D) Exon fragment detection and gene DEFU test for Tregs and memory CD4+/CD25- T cells. (E) Comparison of DEFU genes between Tregs and memory CD4+/CD25- T cells. (F) Comparison of DEFU genes (N=1,112) and quantitatively DS genes (N=403) in Tregs. *** *P*<0.001

### Differences in expression between T1D case and control transcriptomes

Almost all transcripts (Figure 1B) and genes (Figure 1C) in the reduced references were detected (TPM > 0) in both T1D cases and controls. Transcripts that were only detected in cases or only detected in controls were generally of low abundance (Supplementary Figure S1). There were few genes represented in the reduced reference transcriptomes of each cell type that were significantly different between T1D cases and controls (FDR-corrected *P* < 0.05; Figure 1C) in terms of total gene expression. More genes were differentially expressed in memory CD4^+^/CD25^-^T cells (195 of 7,719 genes, 2.5%) than in Tregs (18 of 8,443 genes, 0.2%; Figure 1C).

### Differential splicing in Tregs between T1D cases and controls

Differential splicing was examined between T1D cases and controls, restricting our analysis to only multi-transcript genes defined as those with at least 2 transcripts in the reduced references. 30% of the 8,461 genes in the reduced reference in Tregs were multitranscript genes, as were 28% of 7,733 genes in CD4^+^/CD25^-^ T cells. In memory CD4^+^/CD25^-^ T cells, only 0.3% of multi-transcript genes provided evidence of DS. In contrast, 16% of the multi-transcript genes in Tregs were DS, with significant differences between T1D cases and controls (Figure 1C). The most frequent splicing events in the DS genes in Tregs with the largest changes in Ψ were those consistent with intron retention (Supplementary Table S1).

Differential exon fragment usage (DEFU) between T1D cases and controls was examined to determine if the differential exon usage observed in the 403 DS genes in Tregs extended globally to all genes with exonic expression; and (2) if more DEFU between cases and controls was observed in Tregs than in memory CD4^+^/CD25^-^ T cells. We observed an increased frequency of DEFU (7.1%) in Tregs (1,112 of 15,569 genes; Figure 1D) compared to memory CD4^+^/CD25^-^ T cells (1.2%; 193 of 15,512 genes; Figure 1D). Only 6% of DEFU genes were common to both cell types, consistent with the observed Treg-specificity of differential splicing (Figure 1E). Compared to the set of DS genes in Tregs (Figure 1C), 244 of the 402 (61%) genes with differential isoform usage in Tregs also had significant differential exon usage in Tregs (Figure 1F). Interrogation of unannotated transcriptional events (unannotated junctions and exon-intron border sequences) revealed that in Tregs there was a higher fraction of genes with unannotated events exclusive to T1D cases, also indicative of altered splicing (Supplementary Figure S2).

### Differential splicing alters isoform usage

In addition to determining whether isoforms are expressed at different levels, we also assessed whether the most common isoform changed between cases and controls. Significant changes in transcript rank between T1D cases and controls were more abundant in Tregs than in memory CD4^+^/CD25^-^ T cells (FDR P<0.05; Figure 2). In Tregs, 9% of DS genes also had significant changes in transcript rank (Figure 2A). A majority, 83 of the 113 (73%), of genes with differentially-ranked transcripts had at least one transcript with a rank frequency difference ≥20% between T1D cases and controls, suggesting that the shift in isoform preference is large for those few genes in Tregs with significant changes in transcript rank.

**Figure 2.**
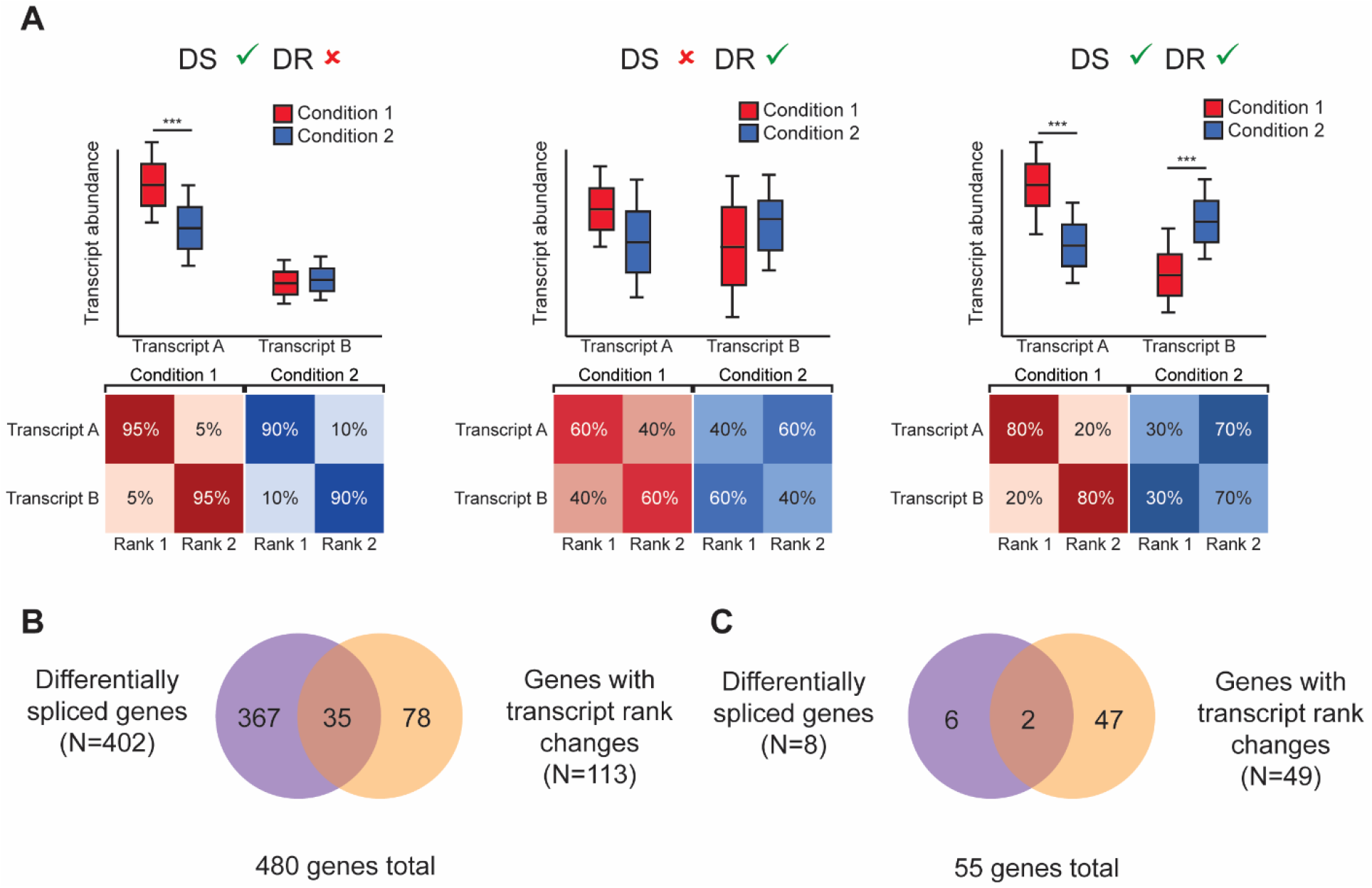
(A) Example of genes with quantitative changes in splicing (differentially spliced, DS) and/or changes in transcript ranking (differentially ranked, DR) between two conditions. Asterisks (***) indicate a hypothetical significant quantitative difference. Summary of transcript rank test and comparison with quantitative DS test for multi-transcript genes in (B) Tregs (N = 2,534 multi-transcript genes) and (C) memory CD4^+^/CD25^-^ T cells (N = 2,184 multi-transcript genes).

In memory CD4^+^/CD25^-^ T cells there were 49 genes with differentially ranked transcripts, and 2 corresponded to genes DS between T1D cases and controls (Figure 2B). Seven of the 49 (33%) genes with significant changes in transcript rank had at least one transcript with a large shift in isoform preference (rank frequency difference ≥20% between T1D cases and controls).

### Differentially spliced genes in Tregs are functionally enriched for RNA recognition motif

We investigated if certain sets of functionally-related or disease-related genes were over-represented among the genes differentially spliced in Tregs. No enrichment for genes located in chromosomal regions associated with T1D (12 of 95 T1D-associated region genes DS; 390 of 2,439 non-T1D-associated region genes DS, *P* = 0.38) or with any autoimmune disorder (56 of 372 autoimmune disease-associated region genes DS; 346 of 2,162 non-autoimmune disease-associated region genes DS, *P* = 0.64) was observed

We tested whether there were any functional protein family motifs that were more likely to be present in proteins encoded by DS genes in Tregs. Genes that encode at least one RNA recognition motif 1 (RRM-1; Pfam accession ID# PF00076.15) were significantly over-represented among DS multi-transcript genes in Tregs, regardless of the splicing test (24/67 (36%) RRM-1-containing multi-transcript genes DS, 378/2467 (15%) other multi-transcript genes DS, FDR-corrected *P*=0.023). Genes containing RRM-1 domains were also more likely to demonstrate DEFU than non-RRM-1 genes in Tregs (50/178 (28%) of RRM-1 genes DEFU; 1,062 /15,391 (7%) of all other genes DEFU; *P*<0.0001).

Because RNA recognition motifs are abundant among splicing factors ^29^, we tested whether splicing factor genes were enriched amongst DS and DEFU genes. Of the 67 known human splicing factor genes from SpliceAid ^30^ mapped to AceView identifiers, 47 were represented in the reduced reference transcriptome for Tregs, and 40 of these were multi-transcript (1.5% of 2,534 multi-transcript genes). Most of these multi-transcript splicing factor genes exhibited differential splicing between T1D cases and controls in Tregs (18/40 (45%) splicing factor genes DS, 384/2,494 (15%) of other multi-transcript genes DS, *P* < 0.0001). No DS splicing factor genes were detected in memory CD4^+^/CD25^-^ T cells. When exons are tested directly without regard to transcript, splicing factor genes were overrepresented in the DEFU genes (33/55 (60%) of splicing factor genes DEFU; 1,079/15,514 (7%) of all other genes DEFU; *P* < 0.0001). This feature was specific to Tregs, as no RRM-1 domain-containing gene or splicing factor gene was considered DEFU in memory CD4^+^/CD25^-^ T cells.

### Features of RRM-1 containing DS splicing factor genes in Tregs

Nine of the 18 multi-transcript splicing factor genes differentially spliced in Tregs are regulated via the differential inclusion of a poison cassette exon (PCE), an exon containing a premature termination codon that is normally skipped, but when included in the mature mRNA can trigger nonsense-mediated decay ^31–33^. These include four members of the serine/arginine-rich splicing factor family (*SRSF5, SRSF7, TRA2A, TRA2B*) and five heterogeneous nuclear ribonucleoproteins (*HNRNPA2B1, HNRNPD, HNRNPH1, HNRNPL* as Aceview gene *HNRNPLandECH1, HNRPDL*). In four cases, it was the differential usage of the PCE that caused these splicing genes to be classified as DS in our analyses –*HNRNPA2B1, SRSF5, SFSF7, TRA2A*, and *TRA2B* (the remaining five genes express PCE-containing transcripts). Thus, differential splicing of these genes can, by altering the availability of key splicing factors, dysregulate the splicing of a much broader set of genes, as we have observed in Tregs. Of note, ten of the dysregulated splicing factors are also either reported as FOXP3 target genes in Tregs (*HNRNPA2B1, HNRNPC, HNRNPD, HNRNPF, HNRNPK, SF1, SFSF5, TRA2A*, and *TRA2B*)^34^ or FOXP3-interacting genes (*HNRNPLandECH1*) ^35^.

### Serine/Arginine-rich Splicing Factor 7 (*SRSF7*)

Among the RRM-1 domain-containing genes with perturbed splicing in T1D, we identified *SRSF7*, a gene that encodes a member of serine/arginine-rich splicing factor gene family and component of the spliceosome ^36–39^. After excluding transcripts with incomplete exon or junction detection, only the *SRSF7*.c and *SRSF7*.l isoforms remained (Figure 3A). Both isoforms are predicted to contain an RRM-1 domain in exon 2 and a C2HC-type zinc knuckle domain in exon 3. The main difference between these two transcripts is the retention of intron 3 in *SRSF7*.l, which introduces a premature stop codon predicted to result in the production of a truncated SRSF7 protein. Part of intron-3 also encodes a well-studied poison cassette exon and the alternative splicing of intron-3 serves as an important regulator of *SRSF7* expression ^36,39–41^. In Tregs from control subjects, the intron-3-retaining *SRSF7*.l contributes little to the total *SRSF7* expression, while the *SRSF7*.c isoform is preferentially expressed and accounts, on average, for 75% of total *SRSF7* expression (Figure 3B). In T1D cases, the expression of *SRSF7*.l increases and contributes ~40% of the total *SRSF7* expression (Figure 3B and 3C), while the expression of the *SRSF7*.c isoform remains on average similar between cases and controls. Total *SRSF7* expression is only marginally higher in cases (Figure 3C). The *SRSF7*.l isoform is frequently the most expressed transcript among T1D cases in Tregs, being preferentially expressed over *SRSF7.c* in 36% of cases compared to 14% in controls. Examination of genomic coverage demonstrates the increase in the retention of intron 3 (Figure 3D, black arrow) without a corresponding decrease the intron-excluding *SRSF7*.c (Figure 3E). Expression of *SRSF7*.c and *SRSF7*.l was similar between T1D cases and controls in memory CD4^+^/CD25^-^ T cells (Figure 3B and 3C), suggesting that T1D-associated changes in *SRSF7* regulation are specific to Tregs. Red line is the average depth for controls; blue line is the average depth for cases; black transcripts are transcripts included in the Treg reduced reference transcriptome; greyed out transcript models are transcripts that were excluded from the reduced transcriptome reference for Tregs. (E) Detailed coverage plot of intron 3 retention showing the mean APN of the intron 3-spanning junction for controls (red) and cases (blue).

**Figure 3.**
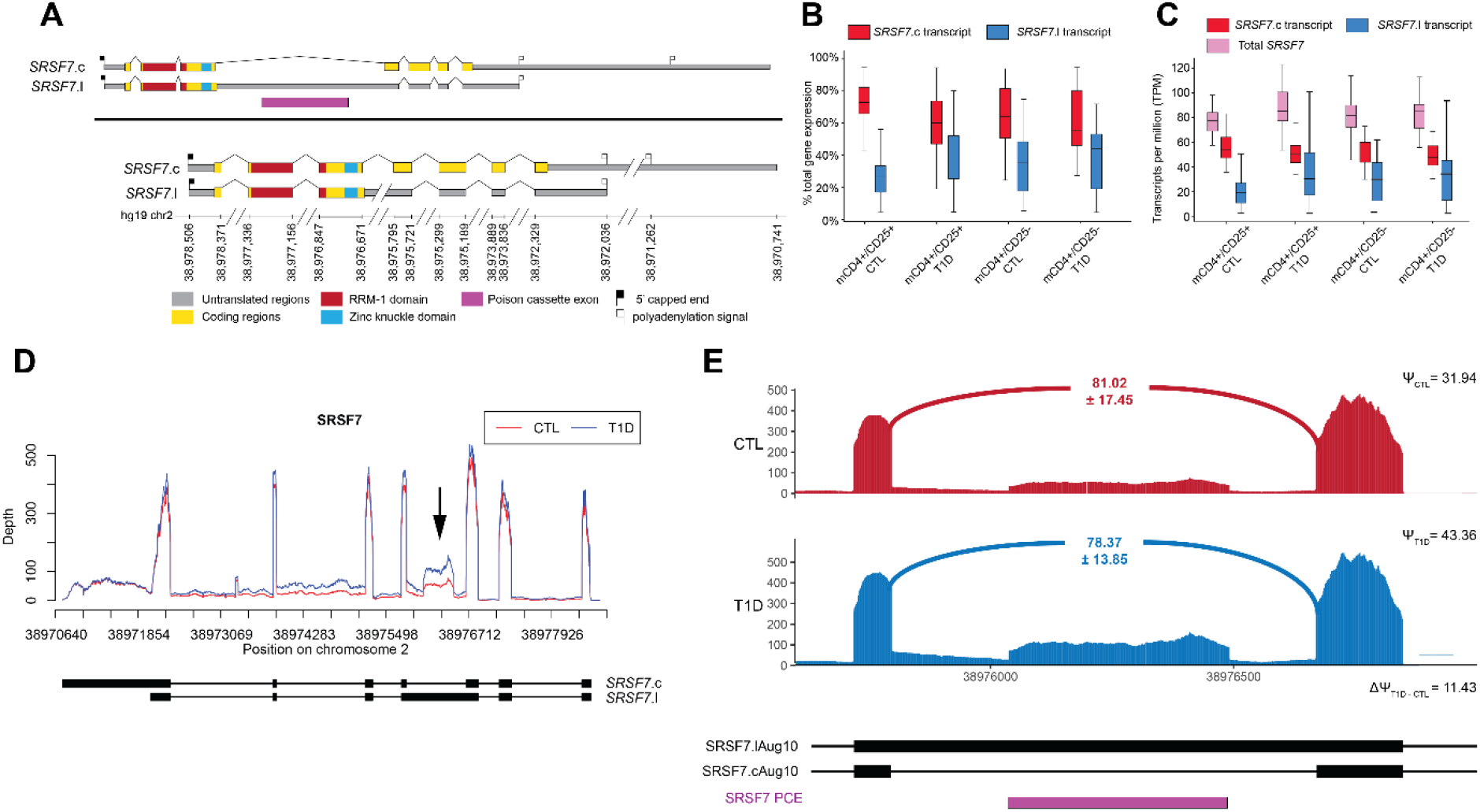
(A) *SRSF7* AceView transcripts represented in the EA-reduced transcriptomes for memory CD4^+^/CD25^+^ T cells and memory CD4+/CD25- T cells. Exonic sequences corresponding to Pfam domains are indicated. Intron and 3’UTR schematics have been reduced in size for visual clarity, as indicated by the broken line marks (“/ /”). (B) Distribution of the proportion of total *SRSF7* gene expression contributed by the *SRSF7*.c and *SRSF7*.l transcripts. (C) Distribution of TPM for total *SRSF7* gene expression and expression of *SRSF7*.c and *SRSF7*0.l transcripts. (D) Genomic coverage plot of *SRSF7* expression in Tregs. Intron 3 is indicated by the black arrow.

### Transformer 2 beta homolog (*TRA2B*)

Seven *TRA2B* transcripts annotated in AceView were included in the reduced transcriptomes for Tregs (Figure 4A), although only two transcripts – *TRA2B*.d and *TRA2B*.g – contain an RRM-1 domain. *TRA2B* is a target of the transcription factor *FOXP3* which is critical for defining Tregs ^34^. Transcripts *TRA2B*.d, *TRA2B*.i, *TRA2B*.j and *TRA2B*.p comprise the bulk of expressed *TRA2B* in Tregs (Figs 4B and 4C). Small increases in the expression of *TRA2B*.j and *TRA2B*.p were seen in Tregs derived from T1D cases compared to controls (Figs 4B and 4C). It has been reported that the second exon of *TRA2B*, which is present in transcripts ‘j’ and ‘p’, acts as a PCE ^31,41^. In Tregs, transcripts retaining this PCE are expressed at higher levels in T1D cases than in controls, while the PCE-skipping transcript ‘d’ is relatively unchanged (Figure 4D, black arrow; Figure 4E). As with *SRSF7*, no such differences in exon/isoform usage were observed in memory CD4^+^/CD25^-^ T cells when comparing T1D cases and controls (Figs 4B and 4C).

**Figure 4.**
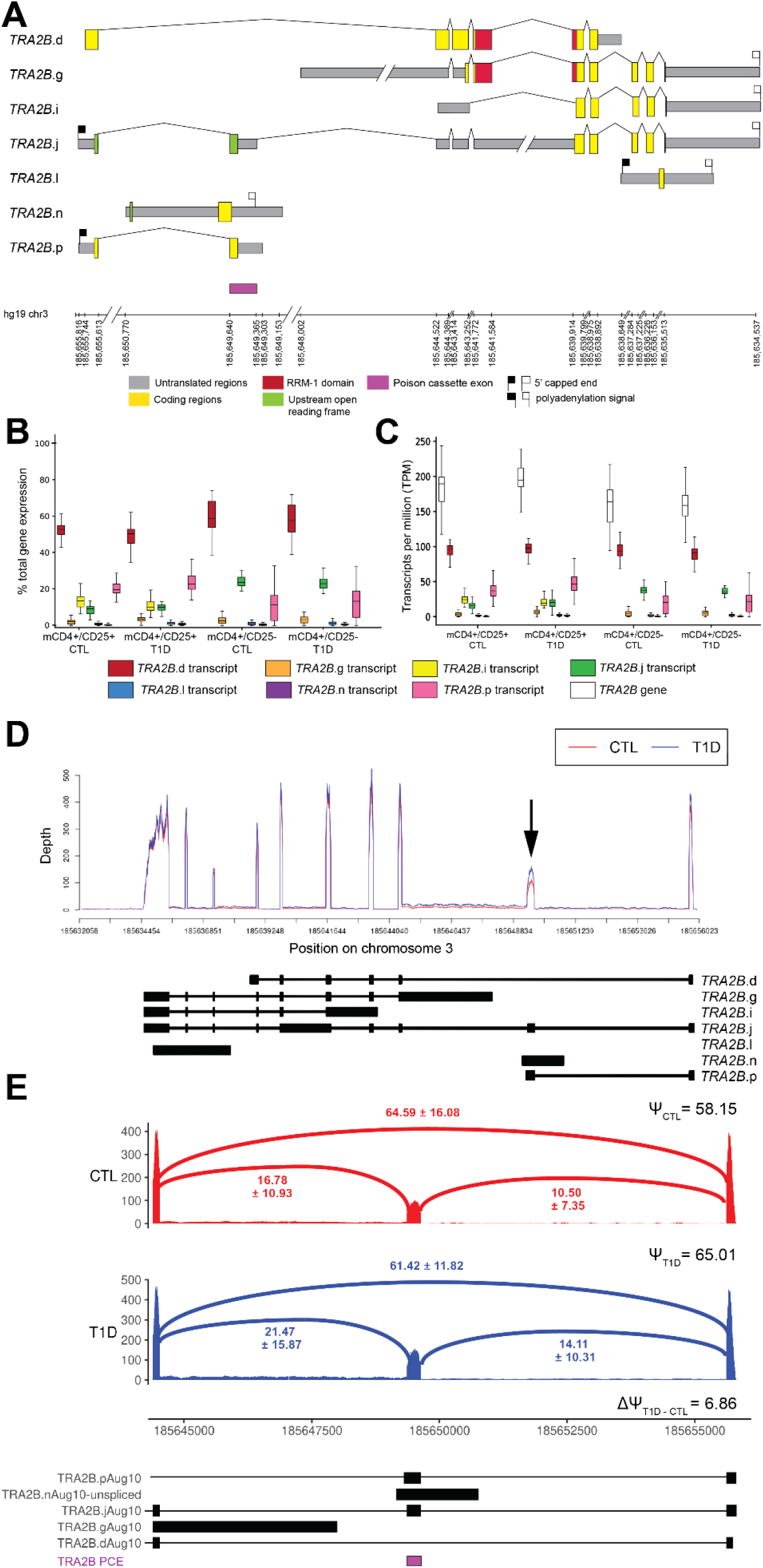
(A) *TRA2B* AceView transcripts represented in the EA-reduced transcriptomes for memory CD4^+^/CD25^+^ T cells and memory CD4+/CD25- T cells. Exonic sequences corresponding to Pfam domains are indicated. Intron and 3’UTR schematics have been reduced in size for visual clarity, as indicated by the broken line marks (“/ /”). (B) Distribution of the proportion of total *TRA2B* gene expression contributed by each expressed transcript. (C) Distribution of TPM for *TRA2B* gene and transcript expression. (D) Genomic coverage plot of *TRA2B* expression in Tregs. The poison cassette exon 2 is indicated by the black arrow. Red line is the average depth for controls; blue line is the average depth for cases; black transcripts are transcripts included in the Treg reduced reference transcriptome. (E) Detailed coverage plot of exon skipping of *TRA2B* exon 2 showing the mean APN of the exon-including and -excluding junctions for controls (red) and cases (blue).

## Discussion

Aberrations in gene regulation via changes in alternative splicing and isoform usage have been implicated in the etiology of many complex disorders ^11–19^. We have previously demonstrated, in a case-only study of T1D, that alternative splicing patterns displayed cell type specificity in lymphocytes, specifically in CD4^+^ T cells, CD8^+^ T cells and CD19^+^ B cells ^26^. There are also several individual reports of changes in isoform production, in lymphocytes, regulated by genetic variants associated with T1D risk ^20–28^. These findings suggest that changes in gene regulation, in the form of alternative splicing, may contribute toward the etiology of T1D in a cell type-specific manner.

Here we explore T1D case-control differences in splicing focusing on a subset of CD4^+^ T cells with particular relevance to autoimmunity, Tregs maintain tolerance to self-antigens by downregulating the induction and proliferation of effector T cells ^42^. Compared to memory CD4^+^/CD25^-^ T cells in which few differences between T1D cases and controls are found, there are several hundred genes exhibiting evidence of differential splicing and isoform usage in Tregs. These alternative splicing events have potentially significant functional consequences and disease relevance as they include many examples of intron retention and are more frequently observed among T1D cases as compared to controls.

Most of the observed examples of intron retention result in the introduction of an in-frame stop codon (79%) and/or a stop codon in all three reading frames ^43^. The retention of these introns is likely to result in the production of truncated proteins with altered or absent function, and/or possibly target the transcript for nonsense mediated decay ^32^. Total gene expression in Tregs is relatively consistent between T1D cases and controls; the primary difference between these subjects is in the ratio of isoform expression of these genes. Gene expression is largely unchanged, but the proportion that each transcript contributes to its gene’s total expression is perturbed in T1D. The prevalence of intron retention and differential exon usage in DS genes suggests that a critical mechanism may be the change in the ratio of functional (e.g. non-intron-retaining) transcripts to non-functional (e.g. intron-retaining) transcripts. Mechanistically, this can be demonstrated by the inclusion of PCEs and other regulatory sequences in mature transcripts in splicing factors and other RNA recognition genes. PCEs are often ultraconserved sequences, suggestive of their importance in gene regulation; in splicing-related genes the role of PCEs is to autoregulate levels of functional protein isoforms ^33,39–41,44,45^. In the *SRSF7* gene, intron 3 contains an in-frame stop codon that is more frequently retained in T1D cases than controls. This intron-retaining transcript contributes more towards the total *SRSF7* gene expression in T1D cases. Alterations in splicing were also identified by variation in transcript rank, resulting in different transcript isoforms being favored in T1D cases compared to controls.

Extending our investigation from transcripts to transcriptional events supported the hypothesis that dysregulated splicing in Tregs is related to the etiology of T1D. Changes in exon fragment usage/expression were observed between T1D cases and controls. We observed an increase in the expression of unannotated transcriptional events likely to impact isoform structure, particularly unannotated junctions (aberrant exon skipping) and exon-intron border junctions (exon-intron read through). In Tregs, more of these events were detected in T1D cases than in controls, indicating the usage of alternative splice sites or simply the failure to efficiently recognize known splice sites. In contrast, there was little difference in the detection of unannotated events in memory CD4^+^/CD25^-^ T cells. The prevalence of intron retention and shifts in preferential isoform usage also suggest that in T1D the regulation of splicing is perturbed in Tregs (but not in memory CD4^+^/CD25^-^ T cells), perhaps through changes in isoform structure.

Several explanations are possible for these observations. Firstly, Tregs may be potentially in a state of cellular stress in T1D, manifesting as malformed (or inefficient) splicing of several genes. The altered splicing that is observed would be incidental to the existing disease state. Second, changes in splicing could be in response to the disease environment of T1D in Tregs. Shifts in isoform usage could serve to regulate gene expression at a post-transcriptional level by diverting proportions of transcripts for a given gene to isoforms that fail to produce fully functional protein. This mechanism could be active even where relatively few differentially expressed genes are observed (as is for Tregs). The abundance of intron retention, the presence of PCEs among DS genes, and the increased expression of unannotated events in cases would seem to support this explanation. Alternatively, transcripts could produce truncated or altered proteins with the potential to disrupt cellular processes by dominant interference. Third, differential splicing in Tregs may reflect Treg phenotypic heterogeneity, or plasticity in their phenotype. Tregs can undergo a phenotypic reversion and a loss of their suppressive functions resulting in a phenotype similar to effector T cells ^46–50^. These ‘ex-Tregs’ have the ability to promote autoimmunity and expansion at sites of inflammation ^46,51–53^. While this is associated with downregulation of FOXP3 ^51,54^, differences in splicing observed in Tregs in T1D could reflect either a higher rate of phenotypic switching or a mechanism that could allow for more rapid switching between phenotypic states. Analysis of Tregs in T1D by single cell RNA-seq may resolve whether the differential splicing observed here is due to Treg heterogeneity in bulk RNA-sequencing data or is cell-intrinsic.

It is possible that the underlying mechanism in T1D may be a combination of these possible alternatives - altered splicing of some critical genes regulating the amount of functional transcript expressed as a response to stress or a consequence of phenotype switching. DS genes in Tregs are enriched for genes that encode RNA-binding proteins and known splicing factors. While splicing factor genes represent only a small fraction of all genes in the genome, ~60% of expressed splicing factor genes have significant changes in isoform usage in Tregs. This suggests that protein-coding genes whose products regulate splicing may be themselves dysregulated in Tregs and consequently their dysregulation propagate and amplify aberrant splicing patterns transcriptome-wide. This could lead to an autocatalytic response where the dysregulation of splicing-related genes leads to aberrant splicing of other splicing genes. The abundance of splicing factors that are differentially spliced in Tregs and that in several instances the splicing difference in question coincides with a reported PCE would suggest that this is a likely explanation for at least some of our observations. Such *trans* regulatory effects, driven by alternative splicing of splicing factor genes and/or other RNA-binding genes may be difficult to specifically delineate given the stochastic nature of transcription and the likely small effect sizes. Most of the shifts in isoform preference in our data are generally small (<20% difference in rank frequency) and are not readily distinguishable from possible *trans* effects. As regulation of alternative splicing may be crucial to proper Treg function ^55,56^, our findings suggest that the preferential expression of alternative transcripts and dysregulated splicing could alter Treg phenotype and function and, ultimately, contribute to T1D risk.

In summary, our findings highlight how alternative splicing and isoform usage can differentiate between T1D-relevant cell types as well as between subjects with and without T1D, even in the absence of significant overall gene expression differences. Our multiple approaches to examining splicing revealed changes in alternative splicing in T1D in Tregs, a critical cell type for maintaining peripheral immune tolerance. Many of the observed differences in isoform usage between T1D cases and controls in Tregs are in the form of changes in isoform structure and frequently involve the mis-splicing of transcripts encoding splicing factors, suggesting possible regulation through an autocatalytic mechanism acting on splicing to alter transcript abundance.

## Methods

### Subject ascertainment

Subjects were ascertained from a study population of 75 T1D cases and 81 age- and sex-matched (non-T1D) controls. The mean age of all participants was 32.6 years (range: 18-49 years), with mean age of T1D onset in cases 19.2 years, with mean duration of disease 13.7 years (Supplementary Table S2). All samples were collected under protocols approved by the Benaroya Research Institute IRB (IRB-07109), with written informed consent obtained from all study participants.

### Sample preparation and RNA sequencing

T cell subsets were purified on an AutoMacs instrument (Miltenyi). Memory CD4^+^ T cells were collected using the Memory CD4^+^ T cell Isolation Kit (Miltenyi) and fractionated into CD25^+^or CD25^-^ subsets by positive selection using on CD25 Microbeads (Miltenyi). Sample purities were assessed weekly during the collection period by flow cytometry (Memory CD4^+^ 95.2%, Memory CD4^+^ CD25^+^ 86.4%, Memory CD4^+^ CD25^-^ 96.6%).

Sufficient RNA for sequencing was purified from 41 controls and 54 T1D cases for memory CD4^+^/CD25^+^ T cells, and 67 controls and 63 T1D cases for memory CD4^+^/CD25^-^ T cells. RNA-seq libraries were prepared according to Illumina protocols and sequenced (2 × 101 nt reads) on an Illumina HiSeq 2000 instrument (137 million ± 58 million reads/sample). Quality of sequencing data was assessed using GC content, and the percentage of adapter content, duplication rate, and homopolymer content in each sample (https://www.bioinformatics.babraham.ac.uk/projects/fastqc/)^57^. Additional quality control procedures to verify sample identity were also applied (Supplementary Methods).

### Annotating transcriptional events

An overview of the approach used to analyze these data is presented in Supplementary Figure S3. We utilized the method of Event Analysis ^58^ to annotate the human transcriptome in terms of exons, exonic regions, individual sequence fragments within these exonic regions based on transcript membership (“exon fragments”), annotated exon-exon junctions, and all other possible, logical exon-exon junctions within a gene. We used AceView gene models for the hg19/GRCh37 genome to assess changes in expression and splicing due to the higher accuracy of its gene models over RefSeq and Ensembl ^59^.

### Quantification of gene expression

Most transcripts in an annotation are unlikely to be expressed; genes not likely expressed in either memory CD4^+^/CD25^+^ Tregs or memory CD4^+^/CD25^-^ T cells were filtered ^58^ as were transcripts not detected in any condition. The remaining transcripts were used to quantify gene and transcript expression. A gene transfer format (GTF) file was generated consisting of only the exons of isoforms included in the reduced reference transcriptome. The program *‘gff_make_annotation’* (https://github.com/yarden/rnaseqlib;^60^) was used to generate a set of annotations for alternative 3’ start sites, alternative 5’ start sites, mutually exclusive exons, skipped exons, and retained introns. Exon fragments and annotated junctions were assigned to each of these annotations as either “inclusion events” (supporting the inclusion of an alternative exon or exonic sequence from a transcript or transcripts) or “exclusion events” (supporting the exclusion of an alternative exon or exonic sequence from a transcript or transcripts). Individual transcriptional events (exonic regions, exon fragments, exon-exon junction, exon-intron border junctions) were quantified ^58^. Sequencing reads were aligned to the generated database of junction reference sequences using Bowtie (version 0.12.9; ^61^) to quantify junction coverage. Reads were aligned to the complete human genome (GRCh37/hg19 version) using the Burrows-Wheeler Aligner for short reads (BWA-MEM, version 0.7.12, ^62^) for coverage of exonic features, namely exonic regions, exon fragments, and introns ^58^. Coverage was calculated for each event as the average depth per nucleotide (APN). An event was considered “detected” if APN ≥ 2 for more than half of all samples per group (cell type × case/control status); i.e., an average of 2 or more mapped reads per feature.

For the analysis of gene expression using transcripts, estimates of transcript abundance were obtained using RNA-seq by Expression-Maximization (RSEM version 1.2.28; ^63^). Cell type-specific reduced transcriptome references were compiled by selecting transcripts with all constituent exons and junctions detected at APN ≥ 2 in either cases or controls ^58^. Transcriptome references were prepared with *‘rsem-prepare-reference’* using the set of transcript sequences as FASTA sequence and a tab-delimited gene-to-transcript file as input ^63^. Default settings were used for all parameters. Transcripts per million (TPM) was the metric used to estimate transcript abundance due to its greater comparability between samples ^64^. A transcript was considered expressed if the TPM > 0 for >50% of all samples per group (cell type × case/control status).

### Differential gene expression and splicing

To assess differential gene expression and splicing in genes represented in the reduced reference transcriptomes, for each gene, the data were modelled as

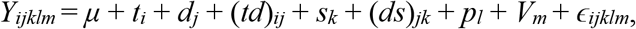

where *Y* is the log-transformed TPM; *t* is transcript *i*; *d* is disease status (*j*=control, case); *td* is the interaction between transcript and disease status; *s* is subject sex (*k*=male, female); ds is the interaction between sex and disease status; *p* is RNA-seq pool (*l* = 1, 2, 3, 4, 5, 6; *i.i.d*. ~N(0,σ^2^_P_)); *V* is a matrix of latent factors for sample *m* used to explain hidden confounders estimated using PEER factors ^65^; and *ε* is the residual (~N(0,σ^2^_P_)). For genes with only a single transcript, this model reduces to

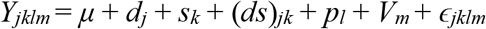

and only differential gene expression is tested for these mono-transcript genes. RNA-seq pool was fit as a random effect to account for the within-pool variance, while all other factors were fit as fixed effects.

If cases preferentially express different transcripts from controls, the *F*-test for the interaction between transcript and disease status (*td*) will be significant, and the gene is considered differentially spliced (DS) ^66^. If there is a difference in the overall expression of a gene, then the *F*-test for disease status (*d*) will be significant, and the gene is considered differentially expressed (DE). Genes that are considered either DS or DE represent the main two hypotheses of interest ^26,67,68^. *P* values were corrected for multiple tests using false discovery rate (FDR) ^69^; a FDR < 0.05 was considered as statistically significant. The model used to assess DE and DS were applied to genes represented by detected exon fragments, with *Y* representing the log-transformed APN; *f* is exon fragment *i;* and *fd* is the interaction between exon fragment and disease status. All analyses of DE and DS were conducted in SAS (v9.4; SAS Institute).

### Identifying splicing differences between transcripts

For each exon with annotated alternative splice variation and for each sample, we averaged the APN of all inclusion events and the APN of all exclusion events. From these estimated a percent spliced in score (PSI, or Ψ) ^60^, was estimated as

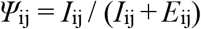

where *I*_ij_ is the average APN of all inclusion events for annotation *i* and subject *j* and E_ij_ is the average APN of all exclusion events for annotation *i* and subject *j*.

A set of annotations was generated to examine the relative expression of alternative first and last exonic regions. The 5’-most (relative to strand) of the first/last exonic regions was considered “reference”; all other first/last exonic regions were classified as “alternative”. For each gene, all possible first and last exonic regions were derived from the set of the transcripts included in the reduced reference. We excluded any first or last exonic region that was also annotated to an internal exon of another transcript, due to ambiguity in defining “reference” and “alternative”. For each annotation and for each sample, we estimated Ψ of the alternative first/last exons, as

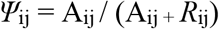

where *A_ij_* is the normalized APN of the alternative exonic region for annotation *i* and subject *j* and R_ij_ is the average normalized APN of reference exonic region for annotation *i* and subject *j*. Splicing differences were annotated in terms of the exonic sequence being included/excluded. For mutually exclusive exons and alternative first and last exons, splicing differences were annotated as the 5’-most exon first, followed by the alternative exon.

### Transcript ranking

A three-tiered ranking system was used to assess changes in isoform preference for genes with two or more transcripts by evaluating whether or not the transcript (or transcripts) that is (are) most/least expressed is (are) changing, without being confounded by small differences in abundance: i.e., sets of transcripts with similar estimates are grouped together. Within each sample and for each gene, transcripts were assigned “rank 1” if their estimated abundance was the highest or within 1 standard deviation of the highest (i.e., the set of <u>M</u>ost <u>E</u>xpressed <u>T</u>ranscripts, METs); “rank 3” transcripts were those with the lowest estimated abundance, or within 1 standard deviation of the lowest (i.e., the set of <u>L</u>east <u>E</u>xpressed <u>T</u>ranscripts, LETs); all other transcripts were assigned “rank 2”. For each condition (cell × status), the frequency of each rank assignment for each transcript was calculated (“rank frequency”).

### Annotation enrichment

Predicted protein family domains from the Pfam database ^70^ for human AceView transcripts were downloaded from the AceView website (ftp://ftp.ncbi.nih.gov/repository/acedb/ncbi_37_Aug10.human.genes/AceView.ncbi_37.pfamhits.txt.gz). Pfam domains were annotated to genes if they were present at least once in the coding region of that gene. Gene set enrichments were performed for all Pfam domains represented in the reduced reference transcriptomes using Fisher’s exact test (JMP Genomics 9, SAS Institute), to identify the function of genes that were significantly more/less likely to be DE or DS compared to other expressed genes. Differences with an FDR-corrected *P* < 0.05 were considered statistically significant.

Enrichment tests targeted specific sets of genes, including RNA Recognition Motif 1 (RRM-1) domain-containing genes, splicing factor genes, autoimmune genes, FoxP3-target genes and FoxP3-interacting genes. Genes containing at least one RRM-1 domain were derived from AceView Pfam annotations using a list of human splicing factor genes obtained from SpliceAid-F ^30^. Human FOXP3 target genes in Tregs were obtained ^34^ as were a list of proteins that interact with human FOXP3 ^35^, and converted to AceView gene identifiers. Genes in chromosomal regions associated with risk for any one of 11 autoimmune diseases were obtained from ImmunoBase (https://genetics.opentargets.org/immunobase). For all tests, a binary response variable was used to indicate gene membership and tested using a χ^2^ test. Statistical significance was considered if the test attains an FDR-corrected *P* < 0.05.

## Data Access

The sequencing data from this study have been submitted to the NCBI database of Genotypes and Phenotypes (dbGaP; https://www.ncbi.nlm.nih.gov/gap; accession number forthcoming.

## Code Availability

All code pertaining to the analysis presented in this study can be found at https://github.com/jrbnewman/T1D_treg_splicing/tree/master

## Supplemental Data

Supplemental Data include 3 figures, 2 tables, and 2 files.

## Acknowledgments

We thank the Benaroya Research Institute (BRI) Center for Interventional Immunology, especially Kassidy Benoscek, Thien-Son Nguyen, Marli McCulloch-Olson, Jani Klein, and McKenzie Lettau, for patient recruitment and sample collection and the BRI Human Immunophenotyping Core for cell isolations. Funding sources: NIDDK/1R01-DK116954 (PC) NIDDK/1DP3-DK085678 (PC and SSR), R01 DK106718 (PC), P01AI042288 (PC).

## Author contributions

Conceived and designed the experiments: SSR, JHB, PC, SO-G; Samples were managed by: SAL, CS, CJG, KC, JHB, PC; Data analysis: JRBN, PC, LMM; Writing - Initial draft: JRBN, PC; Writing – review and edition: all authors

