## Supplementary figures and images for "Shifts in isoform usage underlie transcriptional differences in regulatory T cells in type 1 diabetes"

### Supplementary Figure S1

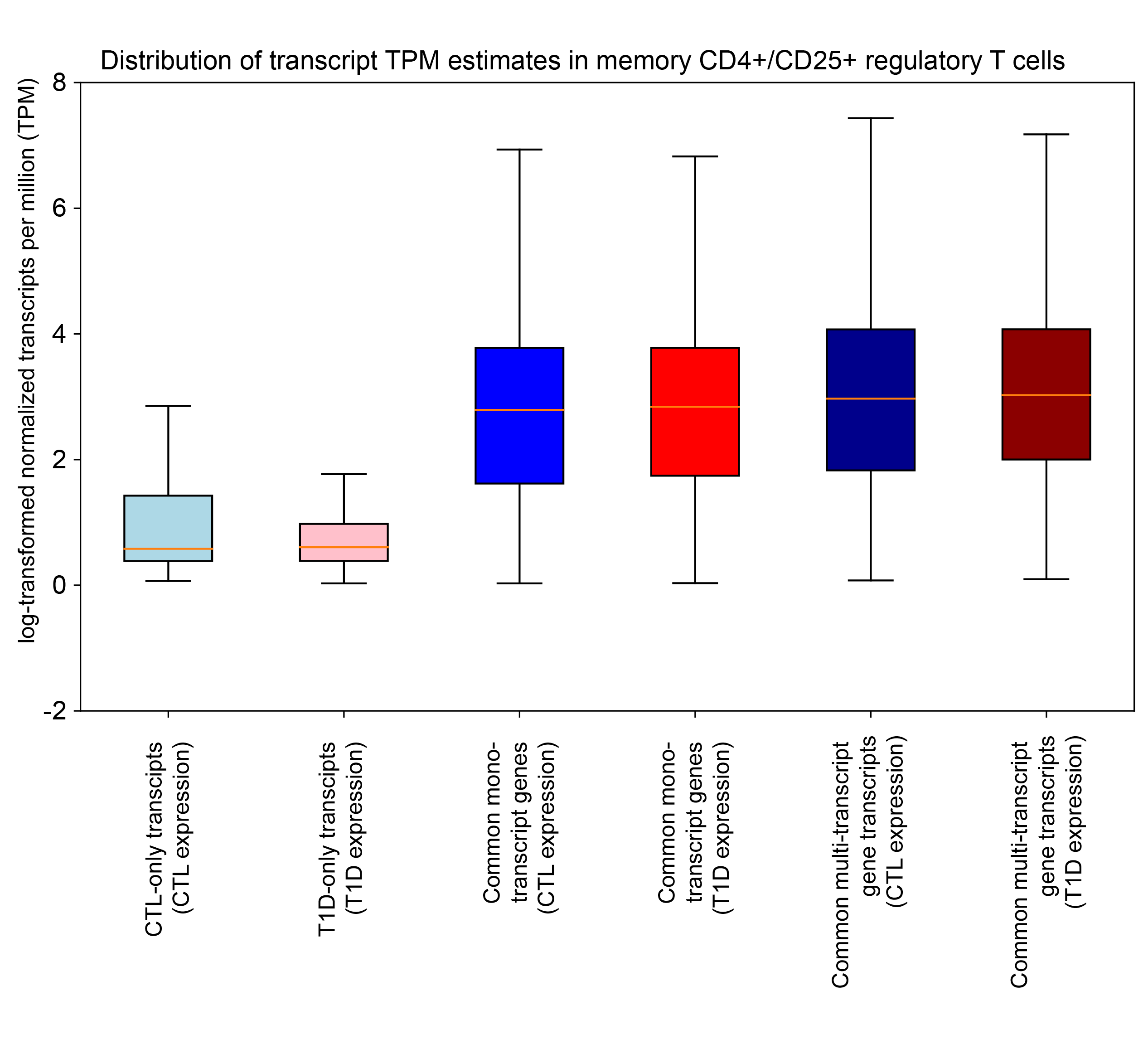

### Supplementary Figure S2

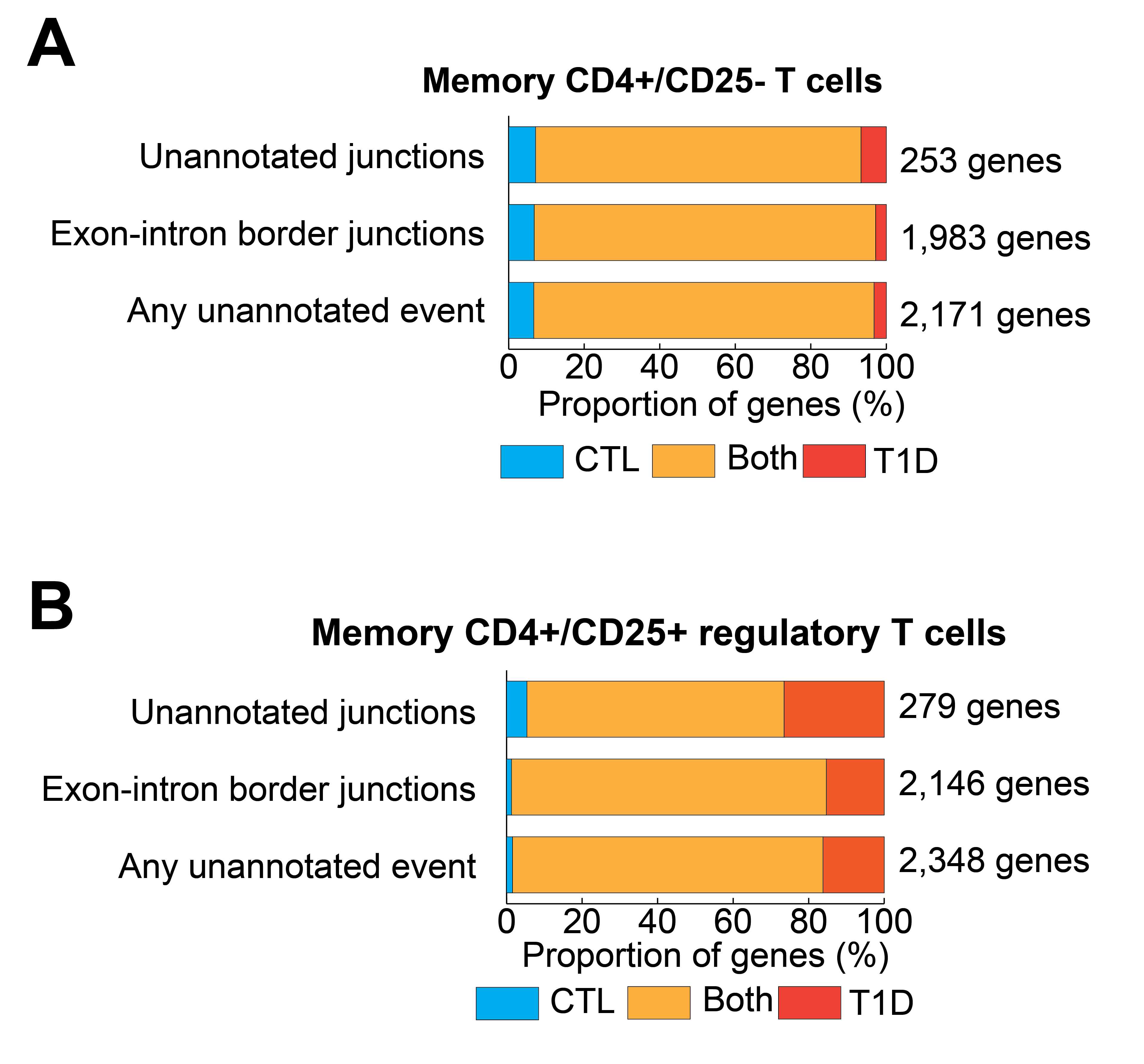

### Supplementary Figure S3

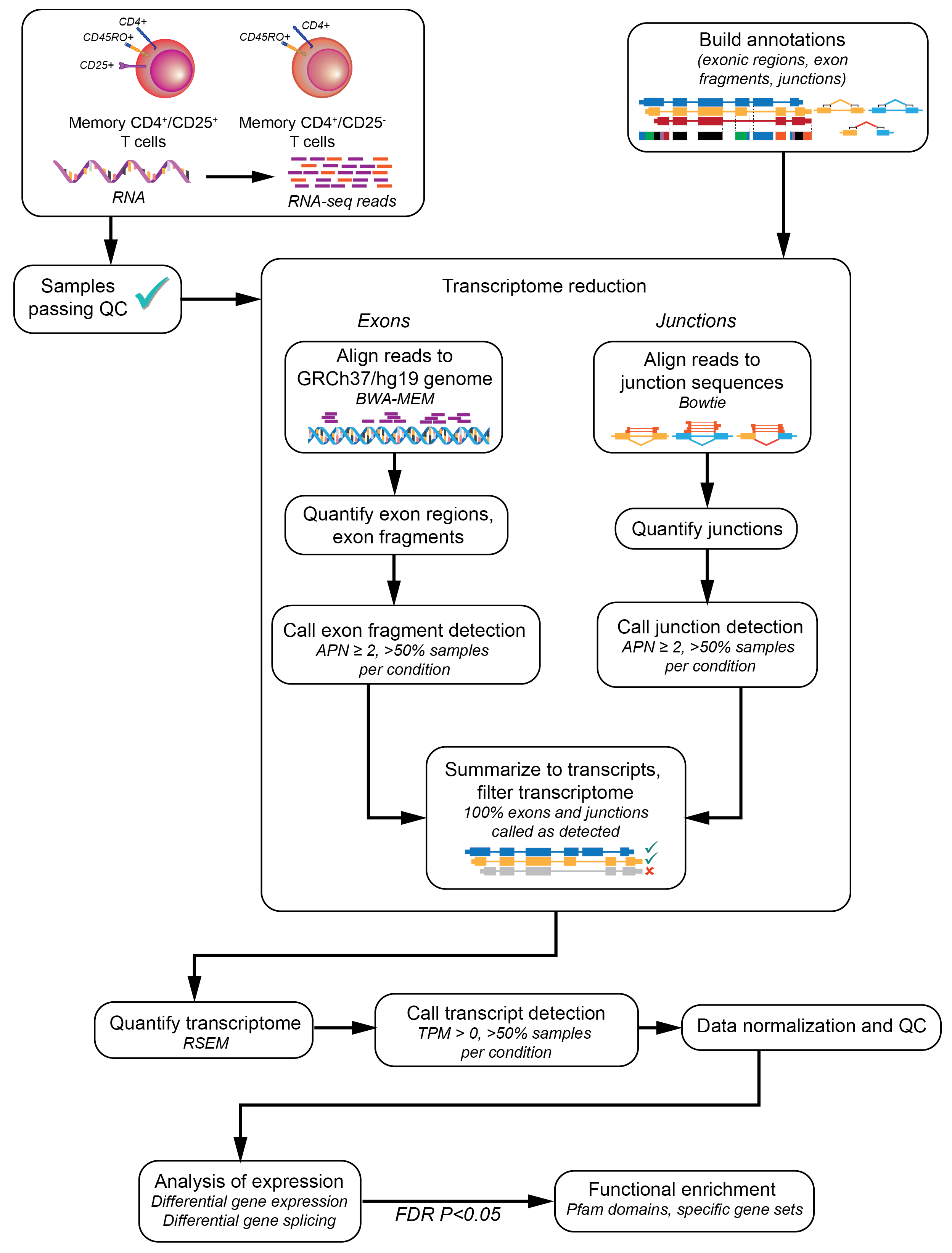
