## Supplementary File S1 for "Shifts in isoform usage underlie transcriptional differences in regulatory T cells in type 1 diabetes"

**Unannotated events in differentially spliced genes**

Given the overrepresentation of splicing or RNA-binding genes and the abundance of intron retention in DS genes in Tregs (Supplementary Table 2), we examined whether novel combinations of existing donor/acceptor sites (unannotated junctions) and/or possible intron read-through (exon-intron border junctions) were more likely to be detected in differentially spliced genes or DEFU genes. Most unannotated transcriptional events (collectively unannotated junctions and exon-intron border junctions) were detected in both T1D cases and controls in memory CD4+/CD25- T cells (86% of genes with unannotated junctions and 90% of genes with border junctions common to both cases and controls; Supplementary Fig 3A). In Tregs, unannotated events were more frequently specific to either T1D cases or controls (68% of genes with unannotated junctions and 83% of genes with border junctions common to both cases and controls; Supplementary Fig 3B). Limiting the analyses to the multi-transcript genes represented in the reduced transcriptome for Tregs, unannotated junctions were more frequently detected in DS genes than in non-DS genes (112/402 DS genes; 429/2,132 non-DS genes; *P*=0.0005). In addition, twice as many DEFU genes expressed unannotated junctions compared with non-DEFU genes in Tregs (226/1,112 DEFU genes (20%), 1,155/11,058 non-DEFU genes (10%); *P*<0.0001), and three times as many genes with unannotated events were exclusive to either T1D cases or controls (108/1,112 DEFU genes with differentially-detected unannotated junctions (10%); 362/11,058 DEFU genes with differentially-detected unannotated junctions (3%); *P*<0.0001). Overall, a higher fraction of genes with unannotated events were exclusive to T1D cases in Tregs (Supplementary Fig 3). Splicing factor genes were also more likely to express unannotated events than other genes in Tregs (13/55 splicing factor genes (24%); 1,368/12,115 other genes (11%); P=0.004).
