## Supplementary Methods for "Shifts in isoform usage underlie transcriptional differences in regulatory T cells in type 1 diabetes"

**Quality control**

To confirm the general similarity of gene expression of samples of the same cell type, normalized expression counts for exonic regions (see Methods, Quantification of gene expression) were analyzed using hierarchical clustering and principal components analysis (JMP Genomics 7, SAS Institute). Expression data were centered and scaled by exonic region to mean of 0 and variance of 1. All parameters were left at their default settings. Samples that did not cluster with their cell type group were either samples of low coverage or otherwise flagged for removal.

To confirm subject sex and donor identity, variant calling was performed from the RNA sequencing data using the Genome Analysis Toolkit (GATK, v3.8.0) (Poplin et al. 2017; Van der Auwera et al. 2013) following the ‘best practices’ for RNAseq short variant discovery. Reads were aligned to the human GRCh37 genome, and duplicate read sequences were marked using Picard ‘MarkDuplicates’ (<https://broadinstitute.github.io/picard/>). Base quality score recalibration was performed using dbSNP release 138, 1000 Genomes Phase 1 SNPs and indels, HapMap release 3.3 SNPs, and Mills and 1000 Genomes gold standard indels as reference variant sites. Genotypes were called running GATK’s ‘HaplotypeCaller’ in GVCF mode (Poplin et al. 2017) and then using ‘GenotypeGVCFs’ as recommended. Variant quality recalibration with a tranche threshold of 99.0% was then applied to minimize false positive calls; as genotype calls were used solely for the purpose of sample identity confirmation, novel variant discovery was not prioritized. Kinship coefficients were estimated using the KING algorithm implemented in PLINK (Chang et al. 2015; Manichaikul et al. 2010) and used to assess genotype concordance between samples from the same individual (e.g. memory CD4^+^/CD25^+^ vs memory CD4^+^/CD25^-^) to confirm their common donor subject. Subjects were also previously genotyping with the ImmunoChip custom genotype array (Illumina) and the Axiom Precision Medicine Research Array (Thermo Fisher) as part of a T1D fine-mapping project (Robertson et al. 2021). Kinship coefficients were estimated between RNA sequencing samples and ImmunoChip samples to assess genotype concordance and confirm sample donor identity.

Chromosome X and Y SNP genotype calls were used to confirm subject sex. In addition, the expression of the genes *TISX* and *XIST* (chromosome X genes involved in X-inactivation) and *EIF1AY* (chromosome Y) was also examined. The ratio of *EIF1AY* to *TISX*/*XIST* expression was calculated, where a high EIF1AY:XIST ratio indicated a male subject and a low or zero ratio indicated a female subject.
