## Supplementary Table S2 for "Shifts in isoform usage underlie transcriptional differences in regulatory T cells in type 1 diabetes"

**Supplementary Table 1. Study sample demographics**

|  | **memory CD4^+^/CD25^+^ regulatory T cells** | | **memory CD4^+^/CD25^-^ T cells** | |
| --- | --- | --- | --- | --- |
|  | **T1D cases** | **Controls** | **T1D cases** | **Controls** |
| Total subjects | 54 | 41 | 63 | 67 |
| Female / Male | 30 / 24 | 20 / 21 | 26 / 37 | 33 / 35 |
| Mean age ± SD | 33 ± 7.6 | 34.3 ± 8.0 | 33.1 ± 8.2 | 32.7 ± 7.7 |
| Mean age at diagnosis ± SD | 18.6 ± 9.2 | n/a | 19.4 ± 10.3 | n/a |
| Mean disease duration ± SD | 14.8 ± 10.6 | n/a | 14.25 ± 10.4 | n/a |
